# Solo: doublet identification via semi-supervised deep learning

**DOI:** 10.1101/841981

**Authors:** Nicholas Bernstein, Nicole Fong, Irene Lam, Margaret Roy, David G. Hendrickson, David R. Kelley

## Abstract

Single cell RNA-seq (scRNA-seq) measurements of gene expression enable an unprecedented high-resolution view into cellular state. However, current methods often result in two or more cells that share the same cell-identifying barcode; these “doublets” violate the fundamental premise of single cell technology and can lead to incorrect inferences. Here, we describe Solo, a semi-supervised deep learning approach that identifies doublets with greater accuracy than existing methods. Solo can be applied in combination with experimental doublet detection methods to further purify scRNA-seq data to true single cells beyond any previous approach.

## 1 Main

Recent advances have enabled transcriptomic and epigenetic profiling at single cell resolution [1]. Single cell genomics can be used to describe the various cell types in a sample, detect changes in cell type composition and gene expression between samples, and track cell lineages and state changes in development and aging [2, 3, 4, 5, 6]. Recent experimental methods that leverage microfluidic droplets or microwells to isolate individual cells have increased the throughput of single cell genomics experiments by an order of magnitude [7, 8, 9]. These protocols isolate single cells by encapsulating them in droplets that contain identifying barcodes. Droplets collect cells according to the Poisson distribution; thus, they encapsulate multiple cells at a rate dependent on the concentration of loaded cells [10, 11]. Instances of multiple cells to a single barcode, referred to as doublets (or multiplets), can depict nonexistent transcriptional profiles that impair and mislead biological inferences from downstream analyses such as low dimensional visualization, clustering, and differential expression (Fig. 1a) [12, 13]. Doublets can be minimized by loading cells at low concentrations, but this greatly increases the cost per cell (Supplementary Fig. 1).

**Figure 1:**
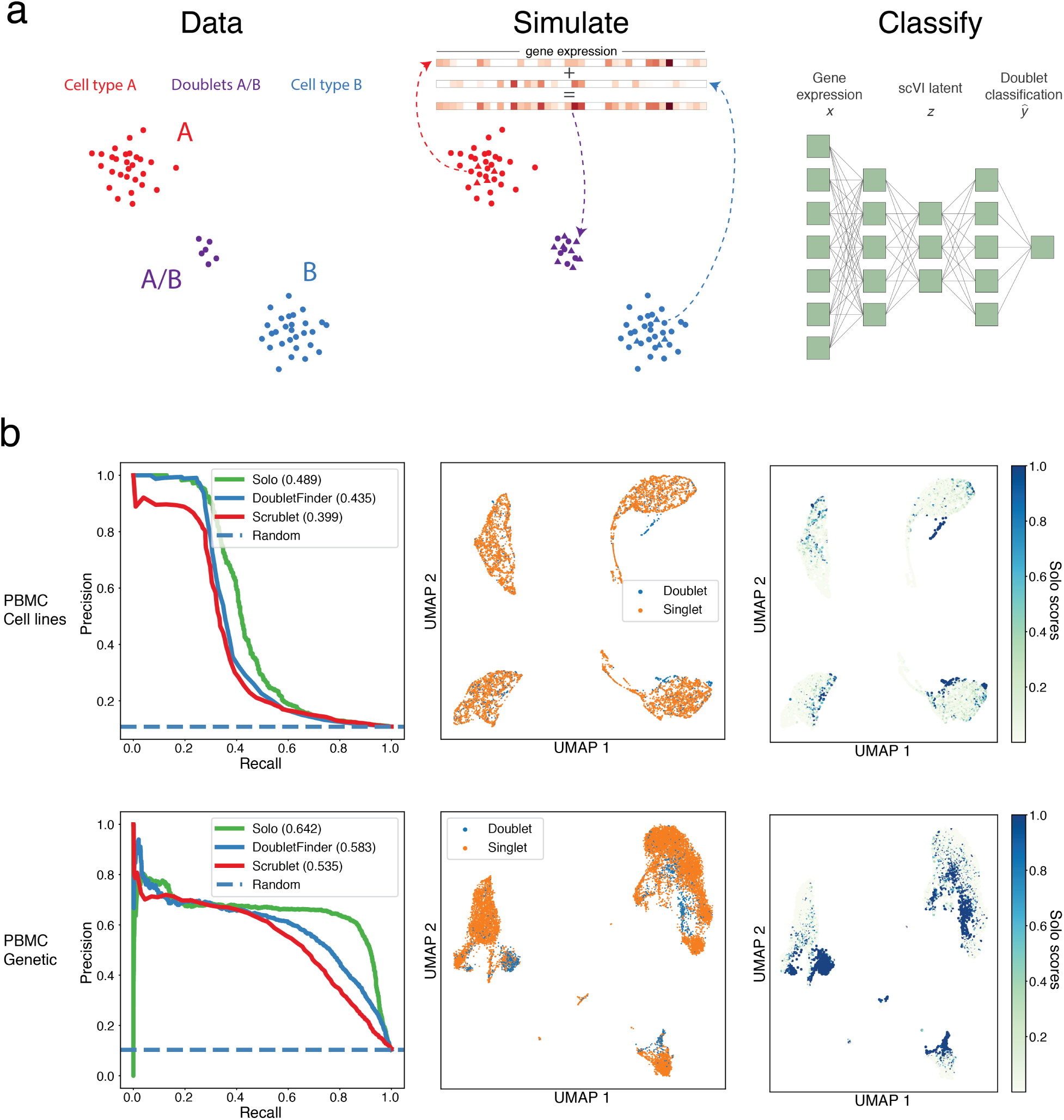
Solo identifies doublets more accurately than previous methods. (a) Doublets combining cells from two distinct types *A* (red) and *B* (blue) create nonexistent and misleading transcriptional states (purple). Assuming that doublets make up a small proportion of the observed data, we can simulate doublets from observed cells to create a training set for a machine learning classifier. Here, we apply a neural network classifier on top of a pre-trained variational autoencoder using the scVI method. (b) We classified doublets using Solo and previous methods Scrublet and DoubletFinder in several experimentally-annotated datasets (Supplementary Tables 1). In the first column, precision-recall curves reveal enhanced doublet identification by Solo. We computed UMAP embeddings of the cells via nearest neighbors in scVI latent variable space. Experimental doublet annotations on the UMAP visualizations in the second column correspond with Solo doublet prediction scores in the third column. In the first row, *Cell lines Antibody* labeled doublets from four distinct cell lines using twelve barcoded antibodies [10]. In the second row, *PBMC Genetic* labeled doublets in unstimulated PBMCs using eight mixed genetic backgrounds and the Demuxlet algorithm [15].

To identify doublets, several groups have proposed experimental protocols to divide cells into subsets and “hash” using an orthogonal information source. The information can come from membrane-bound barcodes [14, 10] or preexisting genetic variation between samples [15] (Supplementary Fig. 2). After sequencing, barcodes with evidence of more than one “hash” identify droplets that contained more than one cell. Although these techniques are effective, genetic variation is unavailable for many experiments and hashing requires additional materials and labor. Alternatively, multiple groups have described machine learning pipelines to classify doublets [12, 13, 16]. These approaches share the following structure: (i) simulate artificial doublets from the profiled cells (mixed singlets and doublets), (ii) train a classifier to identify simulated doublets, and (iii) apply it to the profiled cells to label and remove observed doublets.

Existing computational approaches Scrublet [13] and DoubletFinder [12] operate in linear gene expression embeddings derived by singular value decomposition and apply k-nearest neighbor (k-NN) classifiers. Gene circuits are often nonlinear [17], and gene expression analyses such as batch effect correction, visualization, clustering, and differential expression can benefit from nonlinear embeddings, e.g. those computed using autoencoders [18, 19, 20]. Furthermore, modern neural network classifiers generally perform better than other methods for large datasets [21, 22]. We propose a new pipeline for doublet detection called Solo that employs these enhancements. Solo embeds cells in a nonlinear latent space via a recently introduced variational autoencoder scVI [20] and fits a two layer neural network to classify simulated doublets from observed cells based on these embeddings (Methods, Fig. 1A).

Experimental doublet detection provides labeled cells that can be used to benchmark computational strategies. We studied datasets obtained by cell hashing [10] and genetic variation [15] (Supplementary Table 1). We applied Solo and two recent methods Scrublet [13] and DoubletFinder [12] to several datasets and computed their doublet classification accuracy. Because doublets typically make up 5-10% of cells, we evaluated accuracy primarily with average precision (area under the precision-recall curve, AP) due to its more realistic assessment of imbalanced datasets. Solo classified doublets more accurately than previous methods, boosting AP from second best DoubletFinder’s 0.435 to 0.489 on the hashing dataset and 0.583 to 0.642 on the genetic variation dataset (Fig. 1b) These results were consistent across all experiments performed in these studies (Supplementary Figs. 3, 4; Supplementary Tables 2, 3). Predicted doublets frequently represented nonexistent transcriptional states apparent in 2D visualizations of the cells (Fig. 1b, Supplementary Fig. 3). We asked which components of Solo contribute to its improved accuracy by benchmarking after reducing each component to a baseline version (Supplementary Fig. 5). Both the nonlinear embedding with scVI and NN classifier are critical to optimal performance.

The experimental approaches, cell hashing and genetic variation, have a pre-determined false negative rate that depends on the number of identifying features used, i.e. hash barcodes or genetic sources (Methods). Stoeckius et al. applied both techniques to PBMCs by hashing the separate genetic sources [10]. They observed strong but imperfect agreement between doublet labels based on genetic variation (called by the Demuxlet algorithm) and antibody cell hashing. Demuxlet achieved 0.697 AP classifying doublets with respect to treating the cell hashing labels as ground truth (Supplementary Fig. 6). Solo’s classification accuracy of 0.641 AP approaches that of one experimental method relative to the other. Measured by the area under the ROC curve, Solo’s accuracy exceeds Demuxlet (0.856 to 0.853). Since each genetically distinct sample was tagged with an antibody, the experimental labels are not independent and will tend to simultaneously misclassify doublets that represent different cell types from the same sample. Cells classified as singlets by the experiments and doublets by Solo express more genes than cells classified as singlets by both, indicating that Solo may be identifying many of these expected experimental false negatives (Supplementary Fig. 6).

To explore doublet detection in solid tissue with greater cell type diversity, we performed scRNA-seq on two kidneys from a single mouse using the MULTI-seq hashing method with cholesterol modified oligonucleotides (CMOs) [14]. We analyzed 44,289 cells after basic QC and identified established cell types that have been observed in similar data (Fig. 2; Methods) [23, 24]. In deriving doublet annotations from the CMO counts, existing software proved insufficient because CMO count distributions varied across cell types (Supplementary Fig. 7) [10, 14, 25, 10]. Thus, we developed a novel method, HashSolo, to annotate singlets, doublets, and negative cells from cell hashing counts that better handles this scenario, which frequently occurs in complex samples (e.g. [26]) (Methods). On the jointly profiled PBMC data discussed above, HashSolo’s labels achieve 0.724 AP relative to 0.704 for the authors’ method HTODemux with respect to Demuxlet genetic variation labels (Supplementary Fig. 7, Supplementary Table 4) [10]. For our kidney scRNA-seq, HashSolo rescues thousands of cells labeled as negatives by the existing methods, including the majority of natural killer cells (Supplementary Fig. 7).

**Figure 2:**
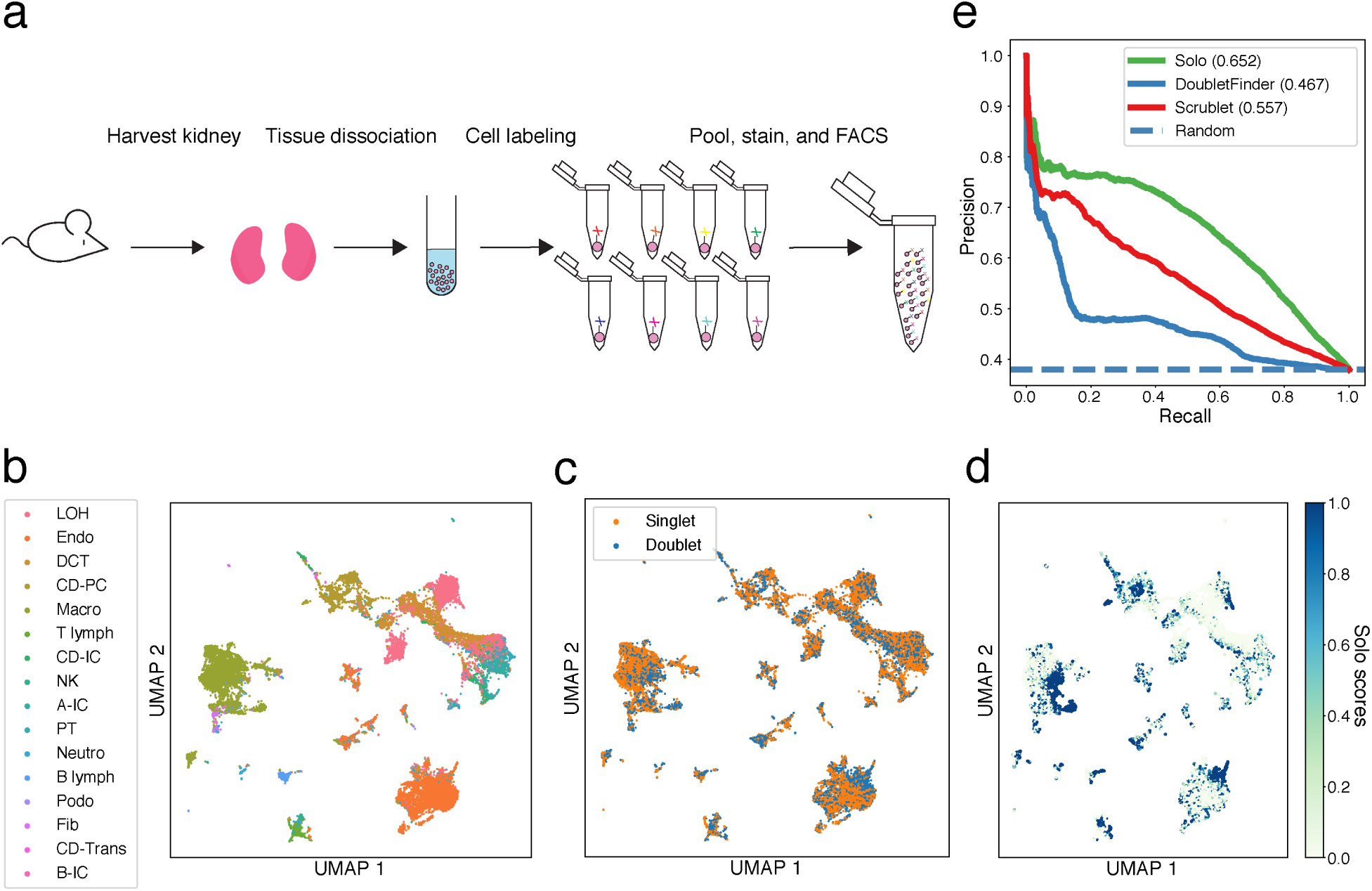
Solo outperforms current methods in kidney tissue with diverse cell types. (a) Schematic overview of samples prior to loading onto 10X Chromium. Dissociated mouse kidney cells were split into 8 tubes and labeled with a unique CMO group, pooled, stained with propidium iodide, and sorted prior to scRNA-seq. (b) We classified the expected cell types, visualized here on a UMAP embedding of the scVI latent space. (c) Doublet annotations derived from experimental CMO hashing tend to exist between and on the edges of clusters. (d) Solo doublet classification scores highlight the same regions. (e) A precision-recall curve shows that Solo classifies CMO doublets more accurately than DoubletFinder and Scrublet on the diverse cell types of the kidney. The legend specifies each method’s average precision.

After establishing CMO-derived doublet labels, we classified doublets using only the transcriptome via Solo. In the kidney, Solo’s accuracy with respect to the CMO labels far exceeds the previous computational doublet identification methods (0.652 AP relative to 0.557 for second best Scrublet) (Fig. 2). Similarly to the other experiments, doublets create nonexistent transcriptional states between and on the edges of cell type clusters that Solo successfully identifies (Supplementary Fig. 8). Solo demonstrated superior doublet classification by either precision-recall or ROC curves on two distinct replicate experiments (Supplementary Fig. 9).

Given the simultaneous advances in both experimental and computational methods for doublet detection, we explored how they can be applied together to achieve even cleaner doublet removal. We created benchmark datasets to simulate a scenario where fewer hashes had been used by merging pairs of hashes; for example, one iteration might merge eight hashes labeled A-H to four hashes AB, CD, EF, and GH (Fig. 3, Supplementary Fig. 10). Solo was able to identify the doublets that were hidden by merging hashes together far beyond random guessing. Thus, we performed additional experiments to tune how the orthogonal predictions were combined to produce a final doublet annotation. We tested linear combinations of the predictions across mixing coefficients (Supplementary Fig. 11). For most datasets, a combined prediction achieved greater accuracy than the experimental or computational predictions alone (Fig. 3, Supplementary Fig. 11). Several attempted pipelines to train Solo on the experimental doublet annotations did not improve results, but more effective joint training schemes may yet exist.

**Figure 3:**
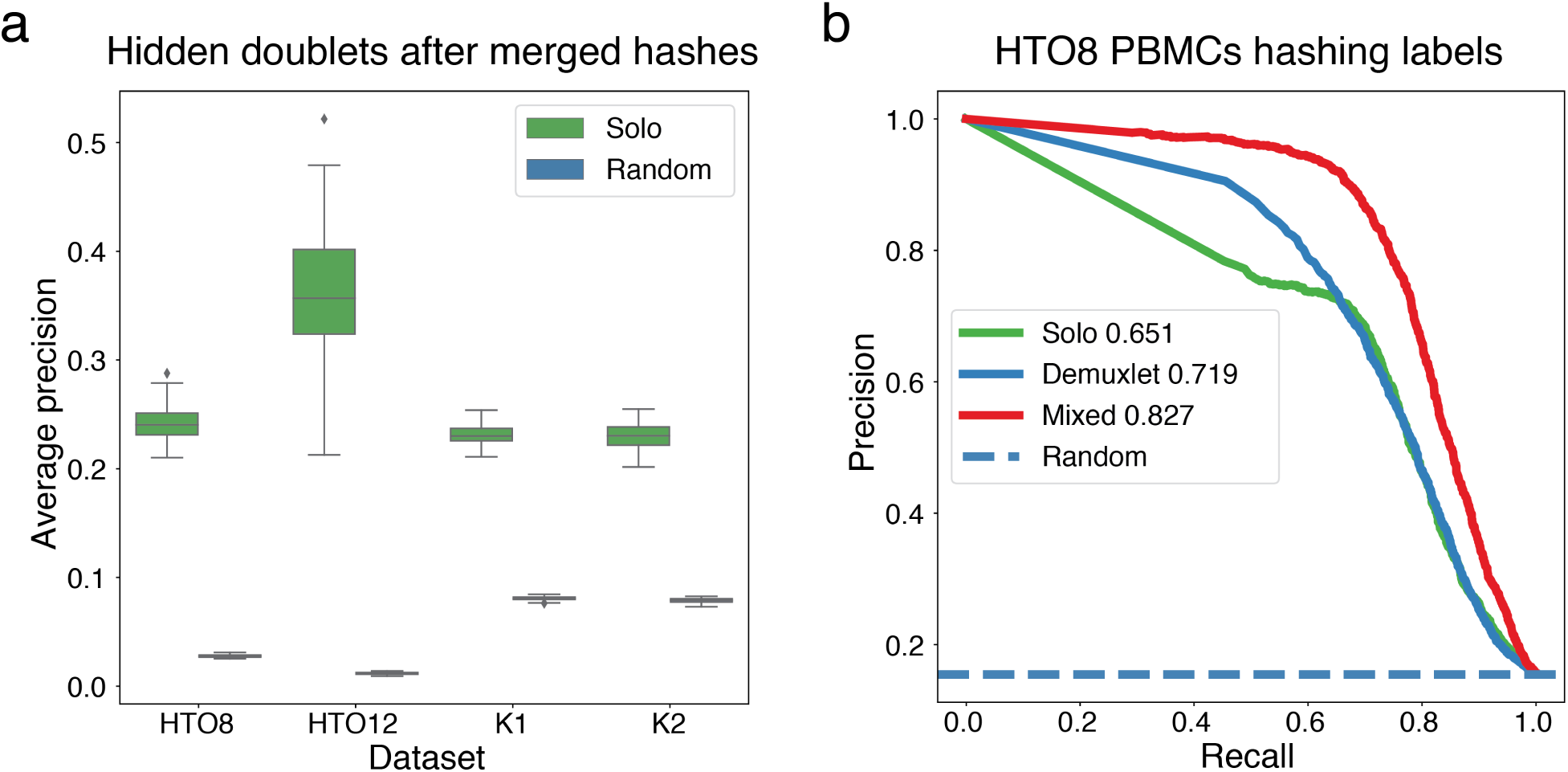
Combining Solo and experimental labels further increases doublet classification accuracy. (a) To assess whether Solo delivers predictive value beyond experimental labels, we simulated datasets by randomly merging pairs of hashes as if half as many hashes has been used (Methods). We then computed accuracy statistics relative to the original hashes on a set of cells that excludes those annotated as doublets by the halved hashes. Even on this challenging set of putative singlets, Solo predictions were accurate beyond random guessing, as measured by average precision. (b) We further considered how to combine Solo and experimental annotations using the Stoeckius et al. HTO8 dataset that was profiled using both antibody conjugated oligos and Demuxlet [10]. Taking the average been Solo and Demuxlet doublet probability calls produces predictions that increase precision-recall over either approach alone with respect to the ACO hashing labels.

In this work, we introduced a novel computational approach for doublet detection called Solo that uses a semi-supervised deep neural network model to represent and classify cells. Solo outperformed existing computational methods for this task on a variety of cell line and tissue datasets with experimental doublet annotations. Computational doublet removal with hashing and Solo reduces costs by enabling high concentration loading in single cell experiments without loss of confidence in downstream analysis and interpretation (Supplementary Fig. 1). Thus, we expect Solo to play a key role in scRNA-seq data processing. Source code is available open source from https://www.github.com/calico/Solo.

## 2 Methods

### 2.1 Solo overview

Computational doublet detection methods assume that most cells in an experiment are singlets and one can estimate a view into the doublet population by generating simulated doublets *in silico* from the observed data. Previous methods and ours operate in three phases: (1) doublet simulation, (2) cell embedding, (3) classifier training.

#### 2.1.1 Doublet simulation

Previous methods simulate doublets by taking the mean or sum of putative singlet profiles [12, 13]. Here, we introduce a procedure to generate a greater diversity of *in silico* doublets. We similarly form the mean profile of two cells, but use it to parameterize a multinomial distribution, from which we sample a count vector. We expose the sampling depth as an option, but default to depth equal to the sum of the two chosen cells. Repeating this procedure, we generate *N*_*d*_ *in silico* doublets. In the machine learning exercise to follow, we fit models to differentiate these *in silico* doublets from the observed data. We chose defaults for doublet depth and number *N*_*d*_ based on test set accuracy on the benchmark datasets (Supplementary Fig. 12).

#### 2.1.2 Cell embedding

Gene expression exists on a low dimension manifold, and standard downstream processing of scRNA-seq exploits this to embed cells with matrix factorization or nonlinear methods. Previous doublet identification methods apply a linear method, PCA to represent cells [12, 13]. Considerable research on scRNA-seq methods has demonstrated significant value from nonlinear methods such as those produced by deep neural networks trained unsupervised as autoencoders [20, 19, 27]. Here, we applied the scVI method to fit a variational autoencoder (VAE) to the observed data [20]. Briefly, VAEs learn a neural network map *q*(*z*|*x*) to encode cell profiles *x* to latent variables *z* and an inverse map *p*(*x*|*z*) to decode the latent variables back to gene expression space [28]. scVI and related single cell autoencoder methods train with stochastic gradient descent to minimize the likelihood of the original count data under a negative binomial parameterized by the decoder. Guided by a parameter sweep across the benchmark datasets (Supplementary Fig. 12), we chose the size of the latent space to be 64 and the number of encoder and decoder hidden units to be 192.

#### 2.1.3 Classification

After embedding the cells in a latent space, previous methods apply variations of a k-NN classifier to predict doublets [12, 13]. Instead, we continue in the neural network framework of scVI and add a standard discriminative classifier to the end of the encoder. More specifically, we fit an scVI model to the observed data unsupervised. Then, we extract the encoder, append an additional hidden layer, and predict doublet status. We freeze the scVI encoder parameters and fit the additional classifier layers to the observed and simulated doublet data via stochastic gradient descent with a binary cross-entropy loss. Continuing to train the encoder weights during doublet classification led to overfitting and decreased generalization accuracy.

The VAE encoder has the added benefit of stochastically embedding the cells to reflect the model’s uncertainty. We found that sampling embeddings from the encoder’s distribution offered regularization that improved generalization accuracy. The scVI model also encodes a library size parameter to reflect the sequencing depth of the cell. Despite prior claims that cell depth is not required for doublet identification [12, 13], we found that passing this feature along to our classifier was critical for effective performance.

#### 2.1.4 Implementation

We implemented Solo in PyTorch as an extension of the scVI software [20]. During training, we performed stochastic gradient descent using the ADAM formulation [29]. We separated 10% of the data as a validation set, evaluated the loss function on the validation set after every epoch, and stopped training once the loss had not improved for 10 epochs. In the hidden neural network layers, we applied dropout regularization with dropout rate 0.2.

### 2.2 Known false negative rates for experimental doublet detection methods

Each experimental doublet detection method uses a known number of identifying features to identify doublets. For cell hashing and MULTI-seq, these identifying features are the number of hashtag barcodes. For Demuxlet, it is the number of genetically diverse individuals. With some probability, doublets will occur from two cells with the same identifying feature and not be appropriately identified. If we assume samples from each identifying feature contribute equally, the rate of unidentified doublets which have the same identifying feature is 1*/f* where *f* is the number of identifying features in an experiment. We refer to 1*/f* as the known experimental false negative rate.

### 2.3 Joint computational and experimental doublet classification

Experimental doublet annotations result in a known false negative rate determined by the number of unique hashes or sources of genetic variation; distinct cells from the same hash or source cannot be identified. In contrast, Solo can identify these doublets via their transcriptional state. To evaluate whether Solo can improve doublet identification on datasets with experimental doublet annotations, we simulated a scenario where we reduced the number of identifying hashes in half by merging pairs of hashes so that many true doublets become undetectable. For example, a set of eight hashing barcodes labeled A through H could be collapsed to four hash barcodes AB, CD, EF, GH. If a cell contains barcodes from A and B, its doublet annotation would be lost for that simulation. We drew random pairs 100 times and quantified Solo accuracy to supplement the remaining available experimental doublet annotations. We then measured how well Solo recovered the lost doublets using average precision.

In addition, we studied PBMC data from eight different patients that was annotated by both cell hashing and Demuxlet by Stoeckius et al. [10]. We applied HashSolo to the experimental annotations to produce a continuous doublet probability score for each technique. We also computed Solo doublet probabilities and explored combining Solo output with one of the experiments while holding the other experiment out to evaluate accuracy. We combined the doublet probabilities as weighted sums across a range of mixing coefficients. As in [10], we restricted Demuxlet doublet labels to those with greater than 95% probability.

### 2.4 Mouse kidney dissociation

A C57Bl/6J mouse was sacrificed and perfused with cold PBS for 6 minutes at a rate of 11 mL per minute. Kidneys were harvested and stored in DMEM on ice. Kidneys were dissociated with 0.2 mg*/*mL of Liberase TM in DMEM at 37°C 800 rpm for 45 minutes (Thermomixer C). Aliquots were taken and visualized under the microscope until tissue was completely dissociated. Liberase TM was inactivated with DMEM+10% fetal bovine serum (FBS) and dissociated cells were filtered through a 100 µm and 40 µm filter. Dissociated cells were then spun down at 200 × *g* for 5 minutes at 4°C and resuspended in PBS+1% BSA.

### 2.5 Cell hashing with cholesterol modified oligos

Anchor/co-anchor cholesterol modified oligonucleotides, sample barcode oligonucleotides, and sample tag library construction were ordered from IDT. Anchor/co-anchor CMOs and sample tag library construction oligo sequences are available from McGinnis et al. [14]. Sample barcode sequences are available in Supplementary Table 5.

Cell concentrations were determined using an automated cell counter (Countess II FL, Invitrogen). 1 million cells were transferred into eight separate 2 mL eppendorf tubes and pelleted at 150 × *g* at 4°C for four minutes. Supernatant was discarded and the pellets were resuspended by gentle finger flicking. Each tube of cells was resuspended in a 90 µL mixture of anchor oligo and three sample barcodes (5 µm final concentration). Samples were gently mixed with light tube tapping and incubated for 5 minutes on ice. 10 µL of co-anchor in PBS was added to each sample (50 µm final concentration). Samples were mixed by light tube tapping and incubated for five minutes on ice. Samples were spun down at 150 × *g* at 4°C for four minutes and washed with PBS+1%BSA. Samples were pelleted at 150 × *g* at 4°C for four minutes and resuspended in 200 µL of PBS with 1 % BSA. Cells were counted on the Countess and the 8 samples were pooled equally. The pooled sample was filtered through a 50 µm filter. Cells were then stained with a viability dye (Propidium Iodide, Sigma P4864 1 µg*/*mL and live cells (PI negative) were sorted using a BD FACSAria (BD Biosciences, San Jose, CA).

### 2.6 scRNA-seq

We loaded a total of 80,000 sorted cells into two wells on the 10x Chromium system (40,000 per well; 10X Genomics). We generated barcoded cDNA libraries using the Chromium Single Cell 3’ Library Construction Kit (V3; 10X Genomics). We constructed sample tag libraries as described in [14]. We quantified final libraries on the Bioanalyzer (Agilent) and sequenced on the HiSeq 4000.

We processed binary base call (BCL) files into cell by gene count and hash count matrixes using BCL2FASTQ (version v2.20.0.422), Kallisto bus (version 0.46.0) [30], and BUStools correct (version 0.39.2) software [31] with default options beyond specifying “-x 10×3” chemistry. We generated our Kallisto reference index from the mm10 Ensembl v96 transcriptome [32].

To ensure cells were of good integrity, we quantified the number of genes, total UMI count, and percent of UMI counts from mitochondrial genes. We removed cells with fewer than 100 genes and 2,000 UMI counts. We removed cells with greater than 60% UMIs coming from mitochondrial genes because they are associated with dying cells. We eliminated genes only expressed in fewer than 20 cells. We performed these analyses and filtering using the Scanpy toolkit [33].

We performed Leiden clustering on a neighborhood graph generated from the first 50 principal components of a PCA on the log counts per million transformation of the data [34]. We annotated cell types using scNym [24] trained on a previously annotated mouse kidney dataset [23].

### 2.7 Hash demultiplexing with HashSolo

After sequencing, the cell by hash barcode counts matrix must be demultiplexed to label each cell with its hash(es) and thus doublet status. In our experiment, we pooled each cell split with three barcodes in order to robustly handle a dysfunctional barcode. Here, we summed the three hashing barcode counts for each sample to obtain sample barcode counts for each cell.

We noticed that existing demultiplexing software for these data—HTODemux [10] and deMULTIseq [14]–were unable to properly classify certain cell types and instead declined to annotate them with a “negative” label. These cells were apparently high quality based on their UMI and unique gene counts and mitochondrial read percentage but did have lower CMO counts.

Therefore, we developed an alternative probabilistic method for demultiplexing cells from hashing counts, which we refer to as HashSolo. Let *h* represent one cell’s hashing count vector, which has length *l* corresponding to the number of distinct hashing barcodes. The hashing count *h*_*i*_ derives from either a signal distribution *θ* if barcode *i* is present for the cell, or a noise distribution *ϕ* if it is not. We modeled these distributions using log-normal.

For each cell, we determined which hash counts will contribute to fitting the signal versus the noise distributions using the following procedure. First, we sorted *h* in ascending order and refer to it as *s*. Then we took *s*_*l*_ as the putative signal barcode count and *s*_1_..*s*_***l***−2_ as putative noise barcodes. We then fit the log-normal parameters of the *θ* and *ϕ* distributions from the putative signal and noise counts across all cells.

For each barcode hashing count vector *h*, we then calculated the likelihood of three different hypotheses that consider the source distribution of the greatest and second greatest counts.

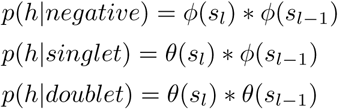

One can set a prior probability for each hypothesis, depending on the dataset of interest. We calculated which hypothesis is most likely using Bayes’ rule.

To be robust to differences between cells with different sequencing depths, HashSolo can be run on homogeneous splits of the full dataset, e.g. by cell type or hash count bins. In our analyses, we either clustered the single cell transcriptomes using the Leiden algorithm [34] or annotated cell types using scNym [24] before running HashSolo for each cluster or cell type.

## 3 Data availability

Mouse kidney CMO scRNA-seq can be accessed through the Gene Expression Omnibus accession GSE140262.

## 4 Acknowledgments

We gratefully acknowledge Jacob Kimmel, Amoolya Singh, and Han Yuan for feedback on the method and manuscript.

## 5 Author contributions

DRK and DGH conceived the project. DRK and NB developed the computational method. NB led the data analysis. NF, IL, MR, and DGH designed the experiment. NF and IL performed the experiment. NB and DRK wrote the manuscript.

## 6 Supplementary Material

**Supplementary Table 1:** Basic details and metrics about the datasets used for benchmarking.

| Dataset | Experimental method | Species | Tissue | Number of cells | Number of doublets | Median UMI count | Median gene count |
| --- | --- | --- | --- | --- | --- | --- | --- |
| Kang et al. Control PBMCs (2c) | Demuxlet | Human | PBMCs | 14619 | 1512 | 1256 | 520 |
| Kang et al. Stimulated PBMCs (2s) | Demuxlet | Human | PBMCs | 14446 | 1552 | 1345 | 546 |
| Kidney pool then FACS 1 (K1) | Cell Hashing (CMOs) | Mouse | Kidney | 24057 | 9131 | 3752 | 1628 |
| Kidney pool then FACS 2 (K2) | Cell Hashing (CMOs) | Mouse | Kidney | 21179 | 7901 | 3929 | 1687 |
| McGinnis et al. (PDX) | MULTI-seq | Human/Mouse | Tumor/Lung | 11697 | 1369 | 2594 | 1087 |
| Stoeckius et al. Cell lines (HTO12) | Cell Hashing (ACOs) | Human | Cell lines | 8191 | 889 | 4636 | 2086 |
| Stoeckius et al. PBMCs (HTO8) | Cell Hashing (ACOs) | Human | PBMCs | 15583 | 2545 | 550 | 321 |

**Supplementary Table 2:**
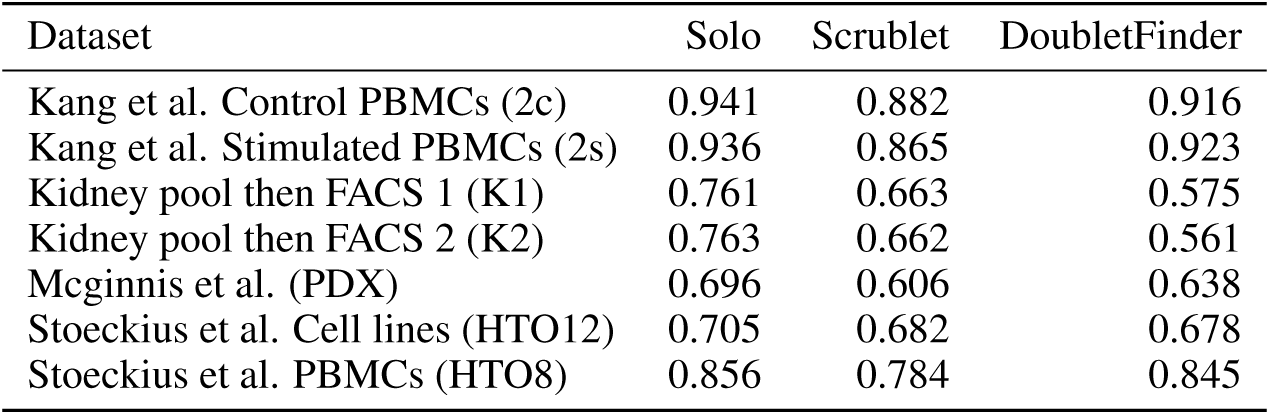
Doublet classification area under the receiver operator curve (AUROC).

**Supplementary Table 3:** Doublet classification average precision (AP).

| Dataset | Solo | Scrublet | DoubletFinder |
| --- | --- | --- | --- |
| Kang et al. Control PBMCs (2c) | 0.627 | 0.535 | 0.583 |
| Kang et al. Stimulated PBMCs (2s) | 0.633 | 0.513 | 0.617 |
| Kidney pool then FACS 1 (K1) | 0.653 | 0.557 | 0.467 |
| Kidney pool then FACS 2 (K2) | 0.650 | 0.548 | 0.450 |
| McGinnis et al. (PDX) | 0.268 | 0.172 | 0.239 |
| Stoeckius et al. Cell lines (HTO12) | 0.486 | 0.399 | 0.435 |
| Stoeckius et al. PBMCs (HTO8) | 0.634 | 0.531 | 0.627 |

**Supplementary Table 4:** Doublet classification via cell hash demultiplexing methods. In the HTO8 dataset, we compared concordance of sample identification between Demuxlet and cell hashing demultiplexing methods.

|  | F1 | Accuracy |
| --- | --- | --- |
| deMULTIseq | 0.8256 | 0.8354 |
| Our method | 0.8261 | 0.8359 |
| Our method with clustering | 0.8246 | 0.8342 |
| htoDemux | 0.8129 | 0.8152 |
| DemuxEM | 0.8039 | 0.8111 |

**Supplementary Table 5:** Sequences of CMO sample barcodes. Template of barcode sequence: 5’-CCTTGGCACCCGAGAATTCCANNNNNNNNNNA30-3’

| CMO Group | Sample Barcode Sequence 1 | Sample Barcode Sequence 2 | Sample Barcode Sequence 3 |
| --- | --- | --- | --- |
| 1 | CGCTCAGTTC | CTACTCAGTC | AACCATTCTC |
| 2 | TATCTGACCT | TCGTCTGACT | GGTTGCCTCT |
| 3 | ATATGAGACG | GAACATACGG | CTAATGATGG |
| 4 | CTTATGGAAT | CCTATGACTC | TCGGCCTATC |
| 5 | TAATCTCGTC | TAATGGCAAG | AGTCAACCAT |
| 6 | GCGCGATGTT | GTGCCGCTTC | GAGCGCAATA |
| 7 | AGAGCACTAG | CGGCAATGGA | AACAAGGCGT |
| 8 | TGCCTTGATC | GCCGTAACCG | GTATGTAGAA |

**Supplementary Figure 1:**
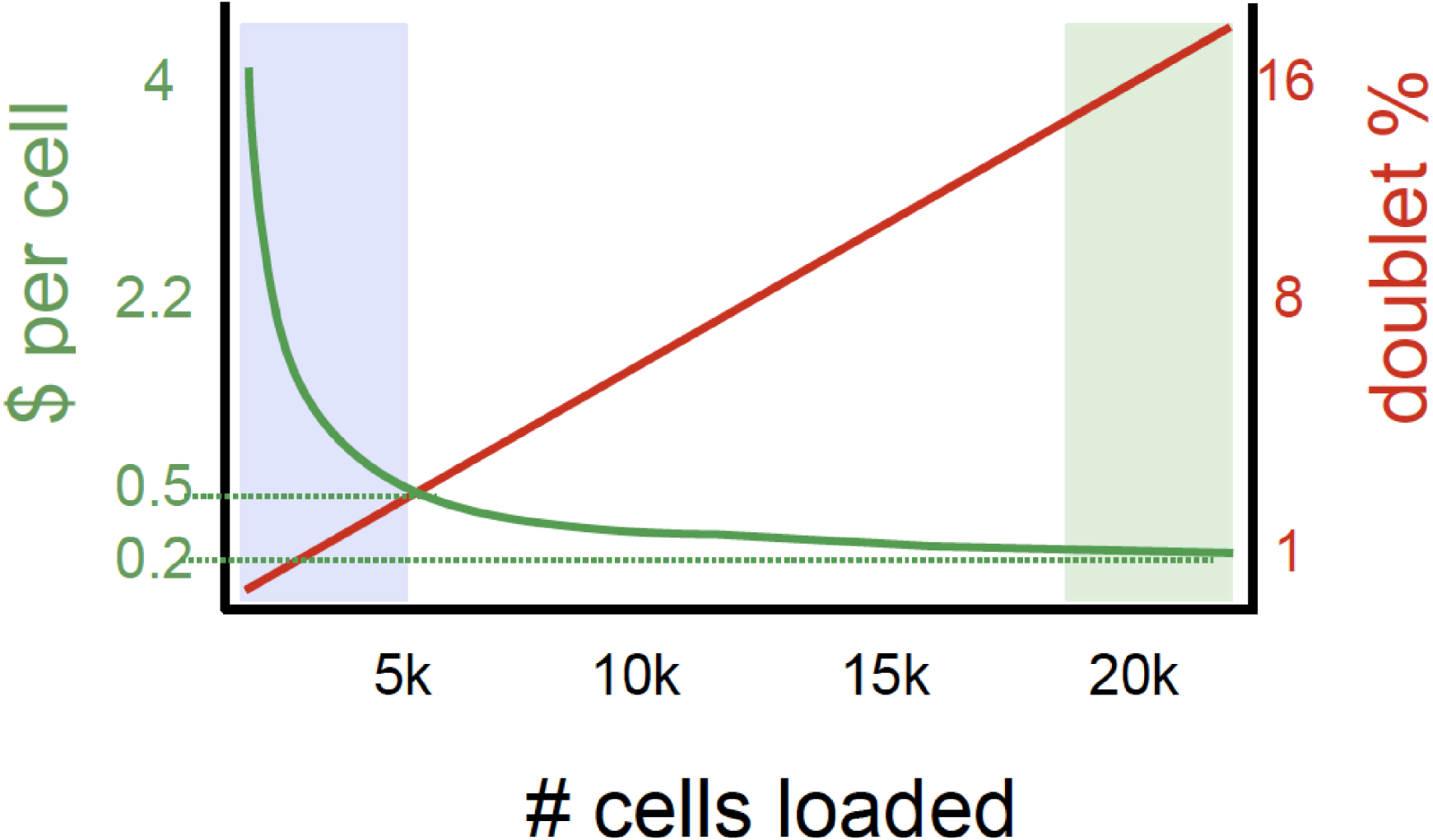
Conservative loading to minimize doublets increases cost per cell. As one loads more cells into droplet and microwell based single cell systems, the cost per cell decreases at the expense of an increased doublet rate. If doublets can be identified and removed, then one can confidently load more cells and achieve lower cost per cell. Cost estimates from https://satijalab.org/costpercell.

**Supplementary Figure 2:**
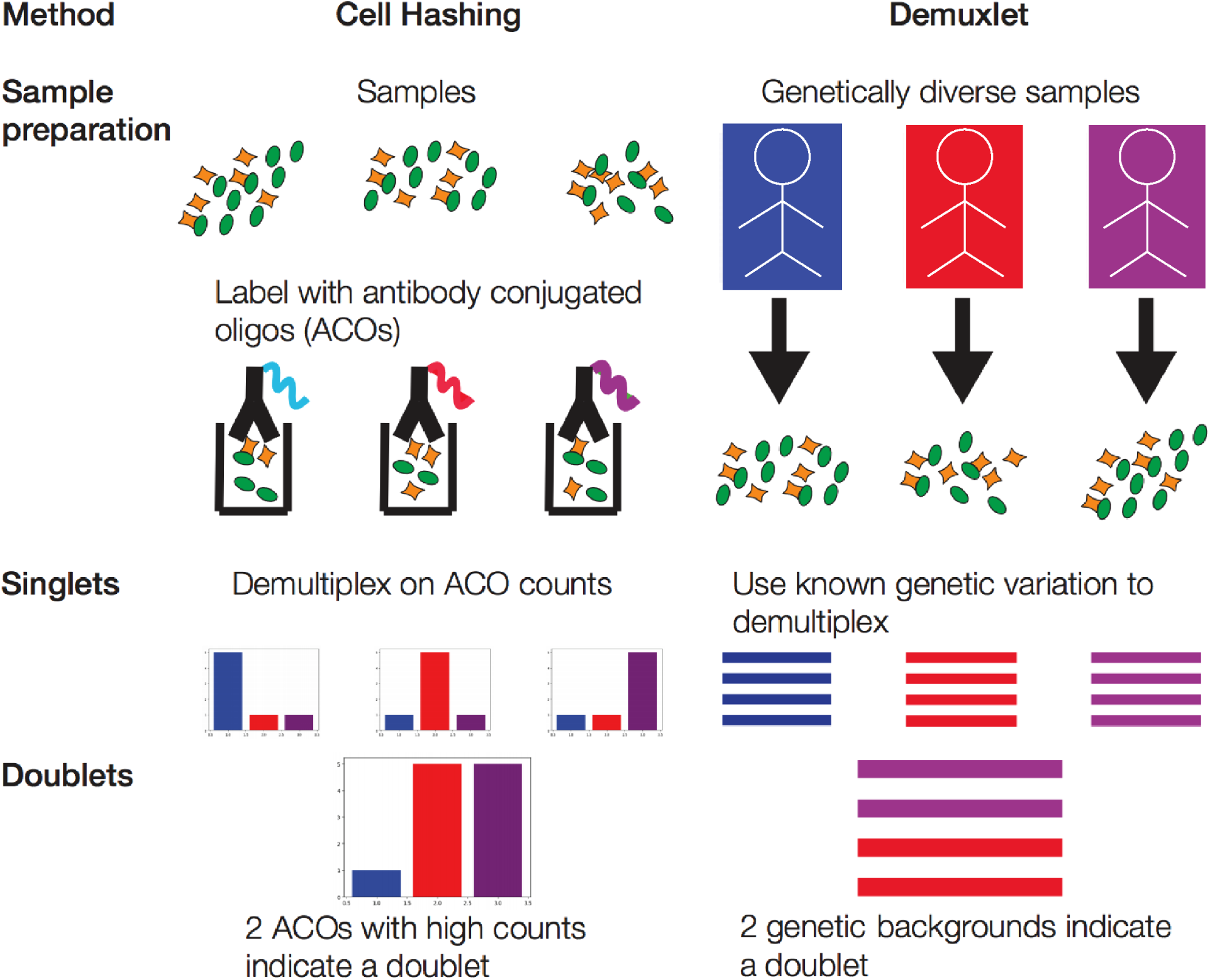
Experimental demultiplexing methods for scRNA-seq. In cell hashing using antibody conjugated oligos (ACOs), (i) dissociated cells are split into a sample for each ACO; (ii) each sample is incubated with an ACO; and (iii) ACO counts for each cell are read out during sequencing. Cells can be associated with their original sample based on their ACO count. Cells with multiple high count ACOs can be recognized as doublets. The Demuxlet approach uses natural genetic variation to identify doublets. Cells from samples with different genetic backgrounds are pooled and encapsulated during the same run. Cells can be associated with their original sample based on observed alleles read from the sequencing reads. Cells with multiple genetic backgrounds can be recognized as doublets.

**Supplementary Figure 3:**
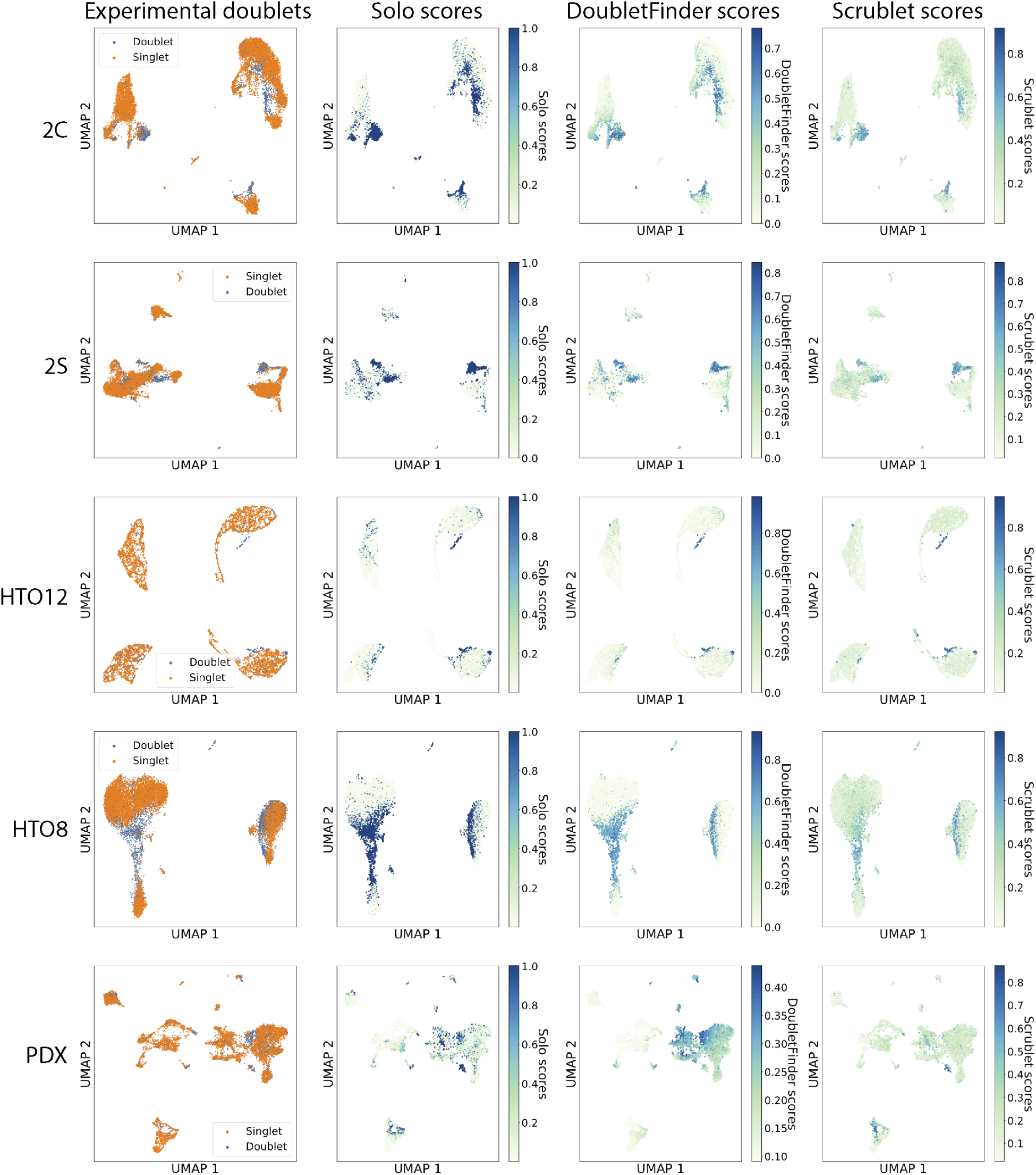
Solo identifies doublets more accurately than previous methods. We classified doublets using Solo, Scrublet, and DoubletFinder in several experimentally-annotated datasets. We derived UMAP embeddings for each dataset from scVI latent variables and colored cells by their (1) Experimental annotation, (2) Solo prediction, (3) DoubletFinder prediction, and (4) Scrublet prediction in each column. The rows represent different datasets: (1) unstimulated PBMCs with genetic variation [15], (2) stimulated PBMCs with genetic variation [15], (3) antibody hashed cell lines [10], (4) antibody hashed PBMCs [10], (5) patient-derived xenograft (PDX) experiment enriched for hCD298+ human tumor cells and mCD45+ mouse immune cells [14].

**Supplementary Figure 4:**
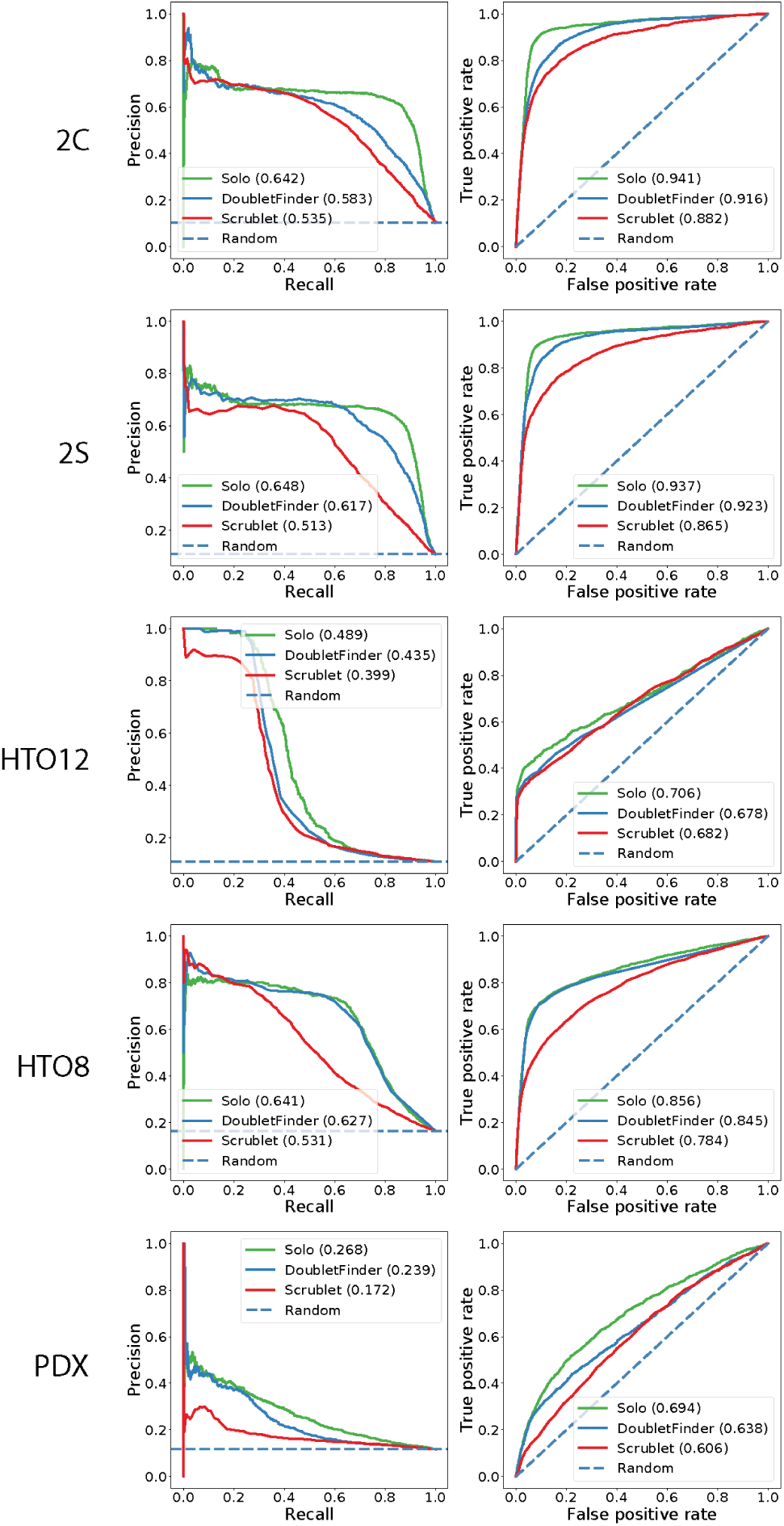
Solo identifies doublets more accurately than previous methods according to AP and AUROC. We classified doublets using Solo, Scrublet, and DoubletFinder in several experimentally-annotated datasets. Precision-recall and ROC curves demonstrate superior predictive ability for Solo. *2s* and *2c* represent stimulated and control PBMCs using Demuxlet [15]. *HTO12* and *HTO8* represent antibody hashed cell line mixture and PBMCs respectively [10]. *PDX* represents hCD298+ human tumor cells and mCD45+ mouse immune cells [14].

**Supplementary Figure 5:**
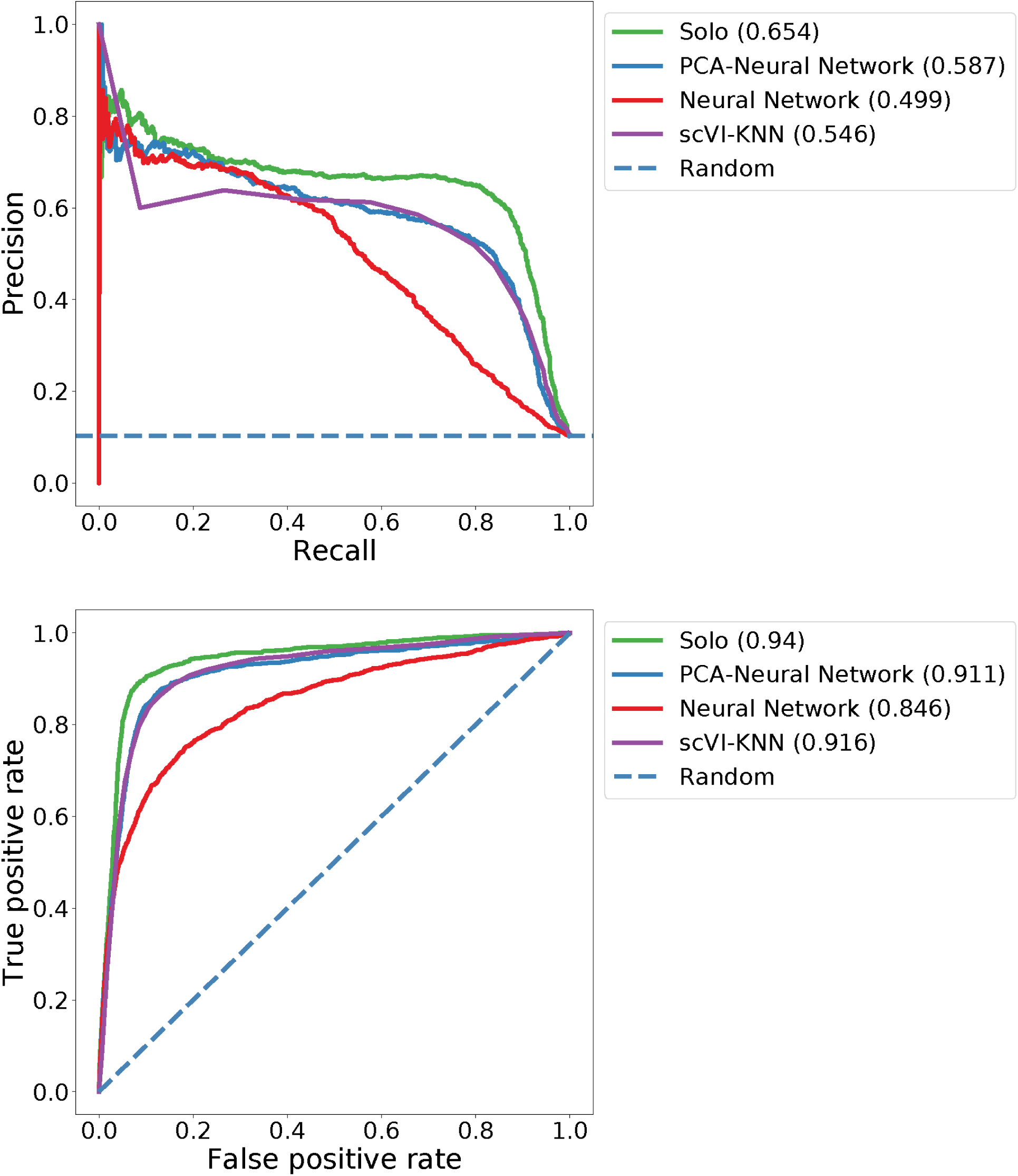
Solo ablation analysis. On the unstimulated PBMCs from [15], we classified doublets and computed accuracy relative to experimental annotations using Solo and several variations where we removed or replaced a step in the method with a baseline version. *PCA-Neural Network* refers to replacing the scVI embedding with PCA. *Neural network* refers to applying a feed forward neural network directly from gene expression without embedding. *scVI-KNN* refers to replacing the neural network classifier with a k-NN classifier on the scVI embedding. Each of these variants achieves lower accuracy than the full Solo method.

**Supplementary Figure 6:**
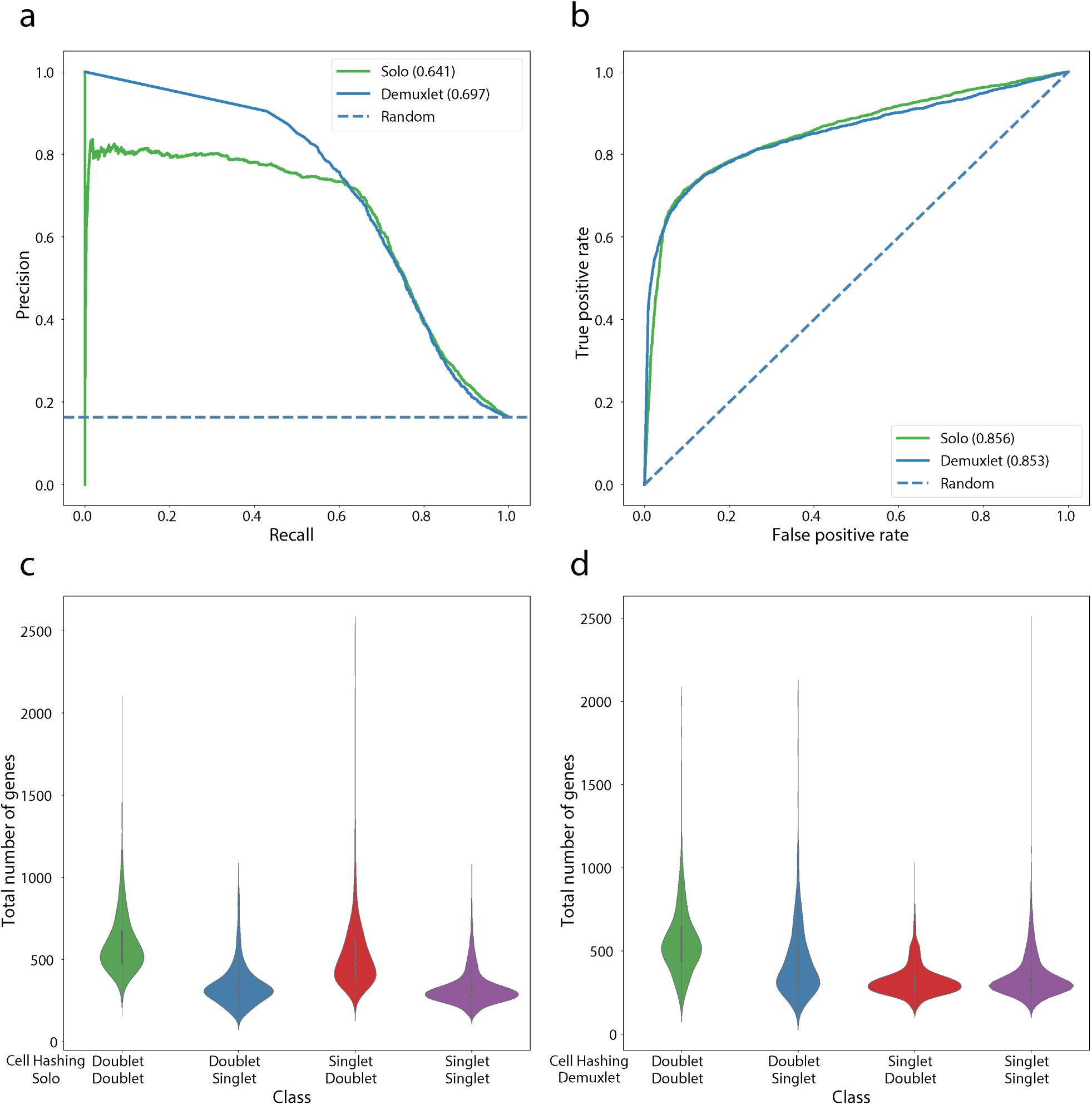
Solo approaches the accuracy of Demuxlet for jointly profiled PBMCs. The Stoeckius et al. HTO8 dataset was generated from PBMCs from eight genetically diverse individuals [10]. Each set of cells was hashed with an antibody conjugated oligo. We compared Solo and the Demuxlet predictions to the hashing labels using precision-recall curves (a) and ROC curves (b). (c) and (d) show the gene count distributions for different prediction classes. Doublets identified by Solo but not cell hashing have transcriptional diversity equivalent to the agreed upon doublets, which suggests that many may be valid.

**Supplementary Figure 7:**
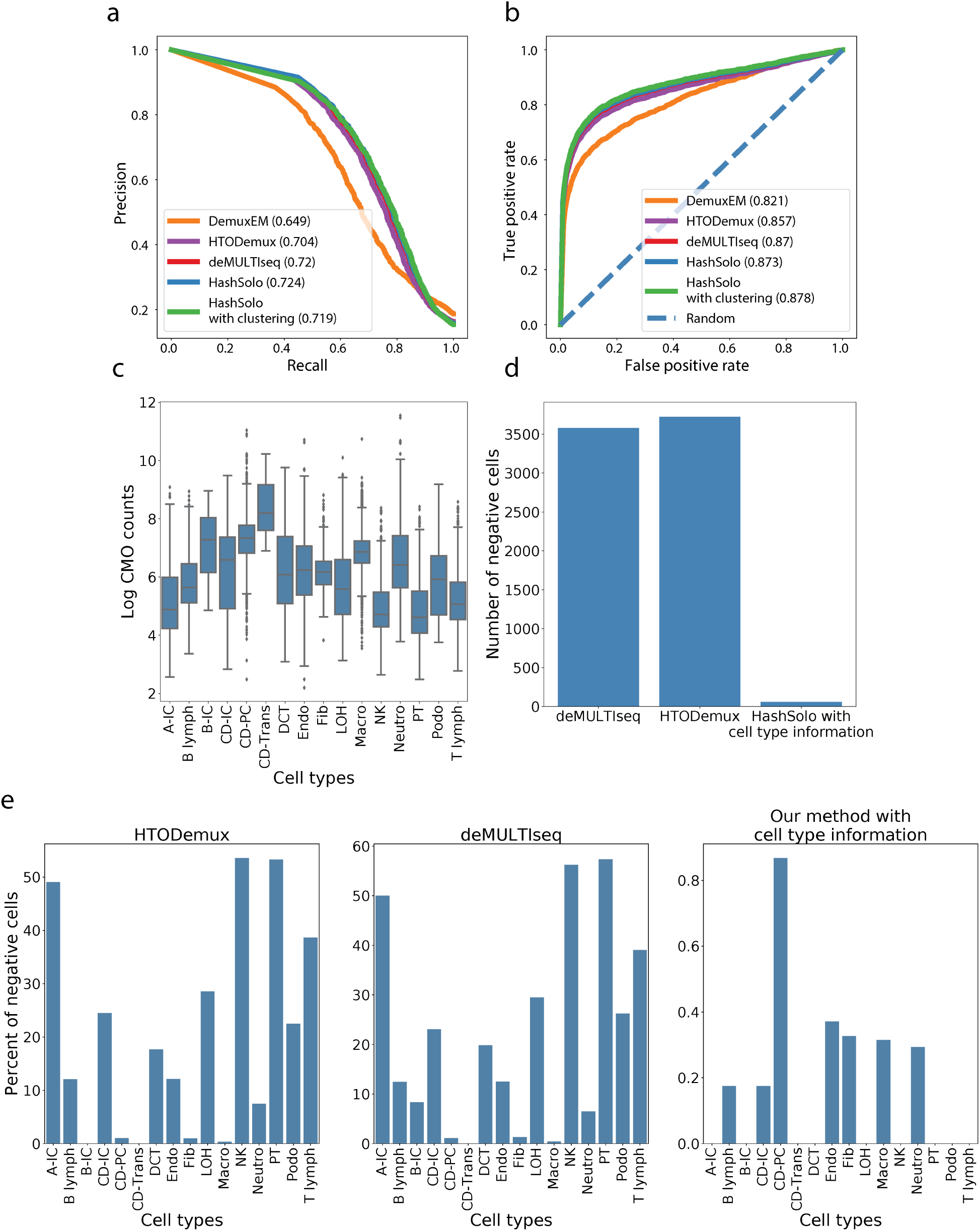
HashSolo outperforms previous methods for demultiplexing cell hashing. The Stoeckius et al. HTO8 dataset was generated from PBMCs from eight genetically diverse individuals [10]. Each set of cells was hashed with an antibody conjugated oligo. We used these data to benchmark hash demultiplexing approaches by comparing doublet annotations to orthogonal annotations derived from the genetic variation and Demuxlet. We ran DemuxEM [25], HTODemux [10], and deMULTIseq [12] according to their recommended settings. For deMULTIseq, we used their semi-supervised method for recovery of negative cells. (a) Precision-recall curves and (b) ROC curves show that HashSolo is a slight improvement upon other methods. (c) CMO count distributions vary across different cell types. (d) HashSolo calls far fewer “negative” cells, in which no singlet/doublet call can be made. (e) HashSolo does not preferentially call negatives for certain cell types with lower hashing counts.

**Supplementary Figure 8:**
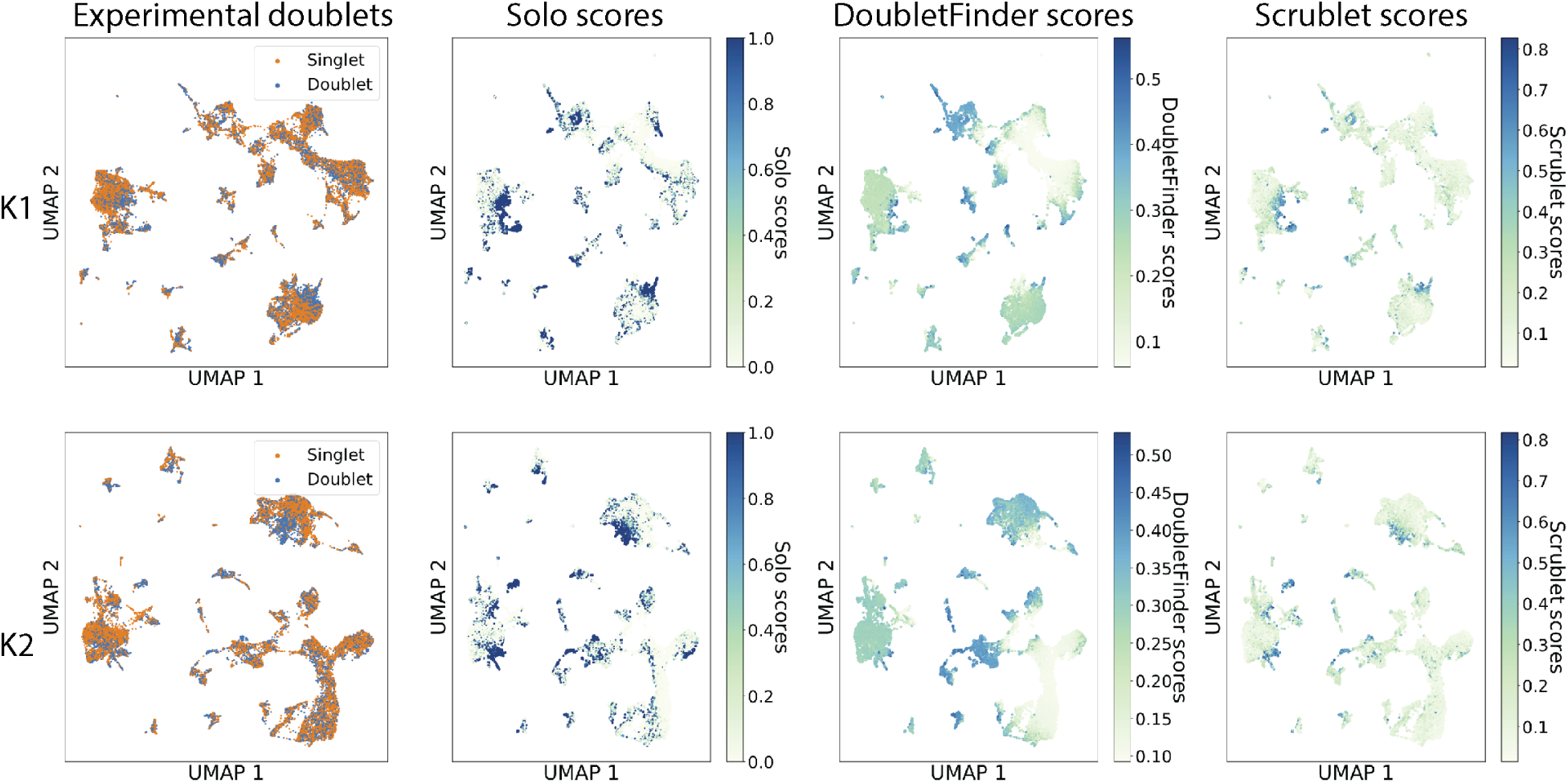
Doublet annotations in CMO-hashed mouse kidney scRNA-seq. We experimentally annotated doublets in two replicate mouse kidney scRNA-seq experiments using cell hashing with CMOs. We derived UMAP embeddings of the cells from scVI latent variables (Methods). We colored cells by doublet predictions for Solo, DoubletFinder, and Scrublet in the right three columns.

**Supplementary Figure 9:**
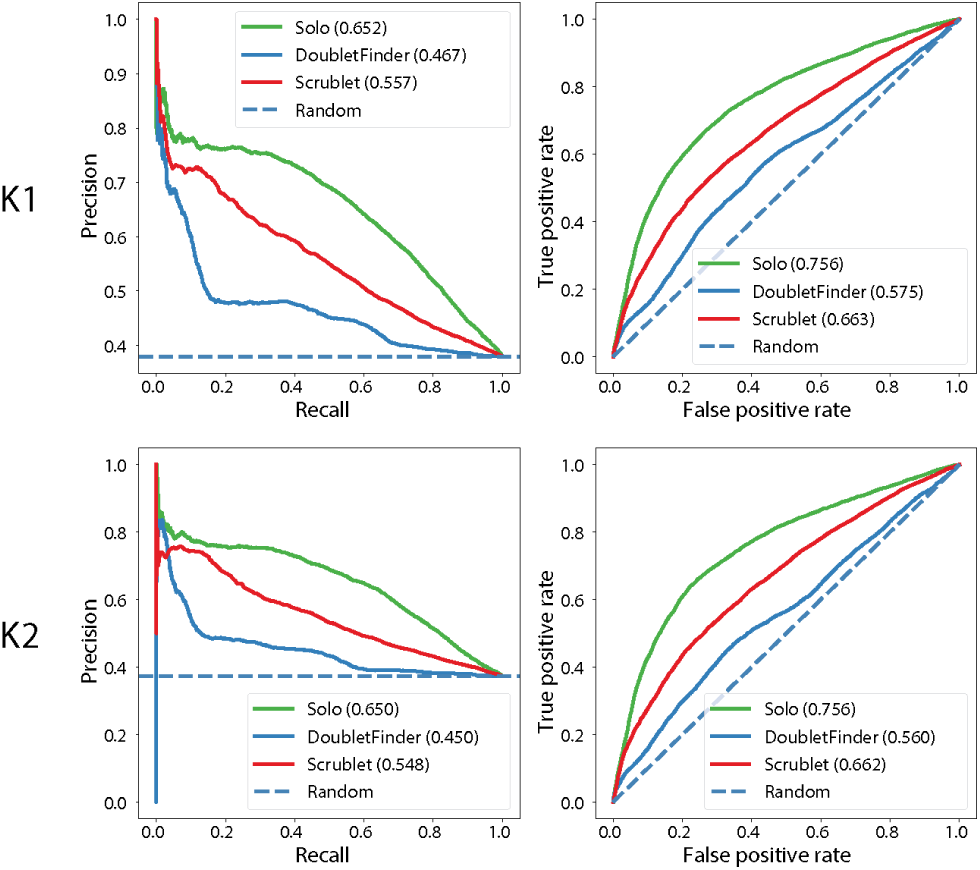
Solo identifies kidney doublets more accurately than previous methods. We experimentally annotated doublets in two replicate mouse kidney scRNA-seq experiments using cell hashing with CMOs. We classified doublets using Solo, DoubletFinder, and Scrublet. Precision-recall and ROC curves demonstrate superior predictive ability for Solo.

**Supplementary Figure 10:**
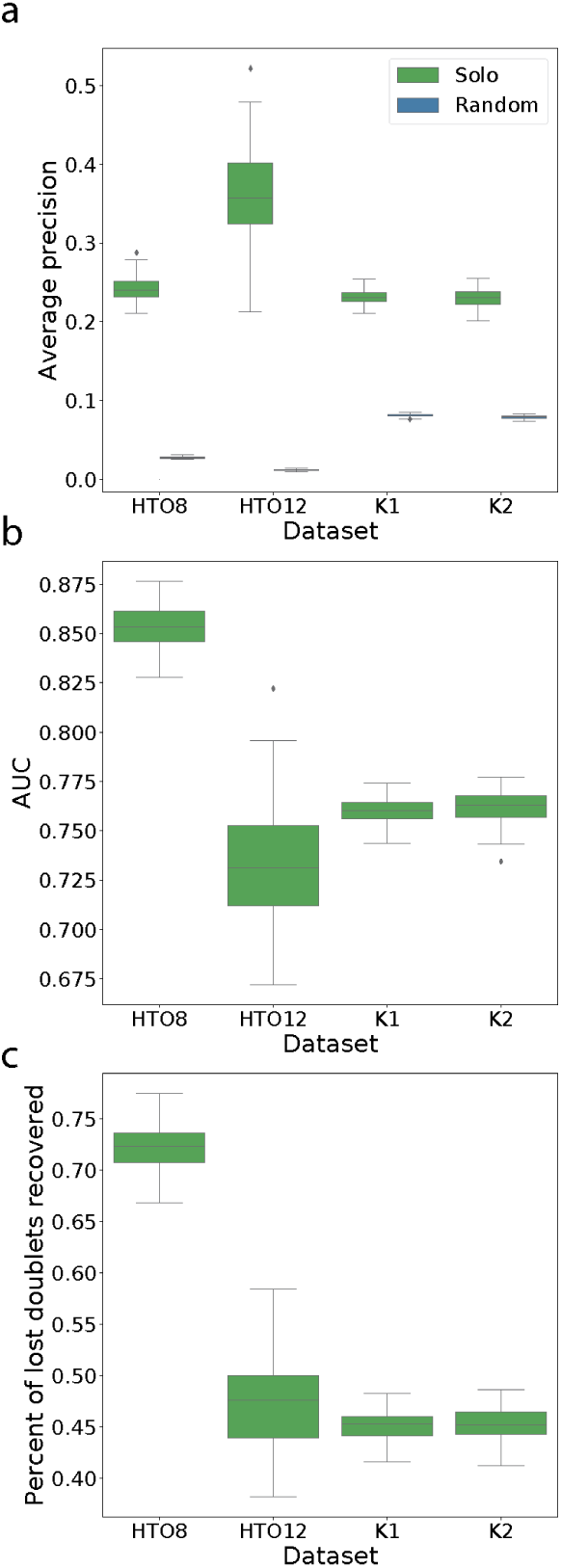
Solo identifies hidden doublets in merged hash experiments. To benchmark a doublet identification strategy that combines an experimental method with Solo, we randomly merged pairs of hashes to simulate having used half as many hashes (Methods). In these experiments, we computed accuracy statistics on a set of cells that excludes those annotated as doublets by the halved hashes. Thus, we quantify the ability of Solo to identify the remaining unlabeled doublets, a more challenging task than the standard setup earlier in the paper. (a,b) Solo AP and AUROC for 100 iterations of randomly merged hashes are always accurate beyond random guessing. (c) Solo recovers 45-75% of doublets missed by the halved hashes in these simulations.

**Supplementary Figure 11:**
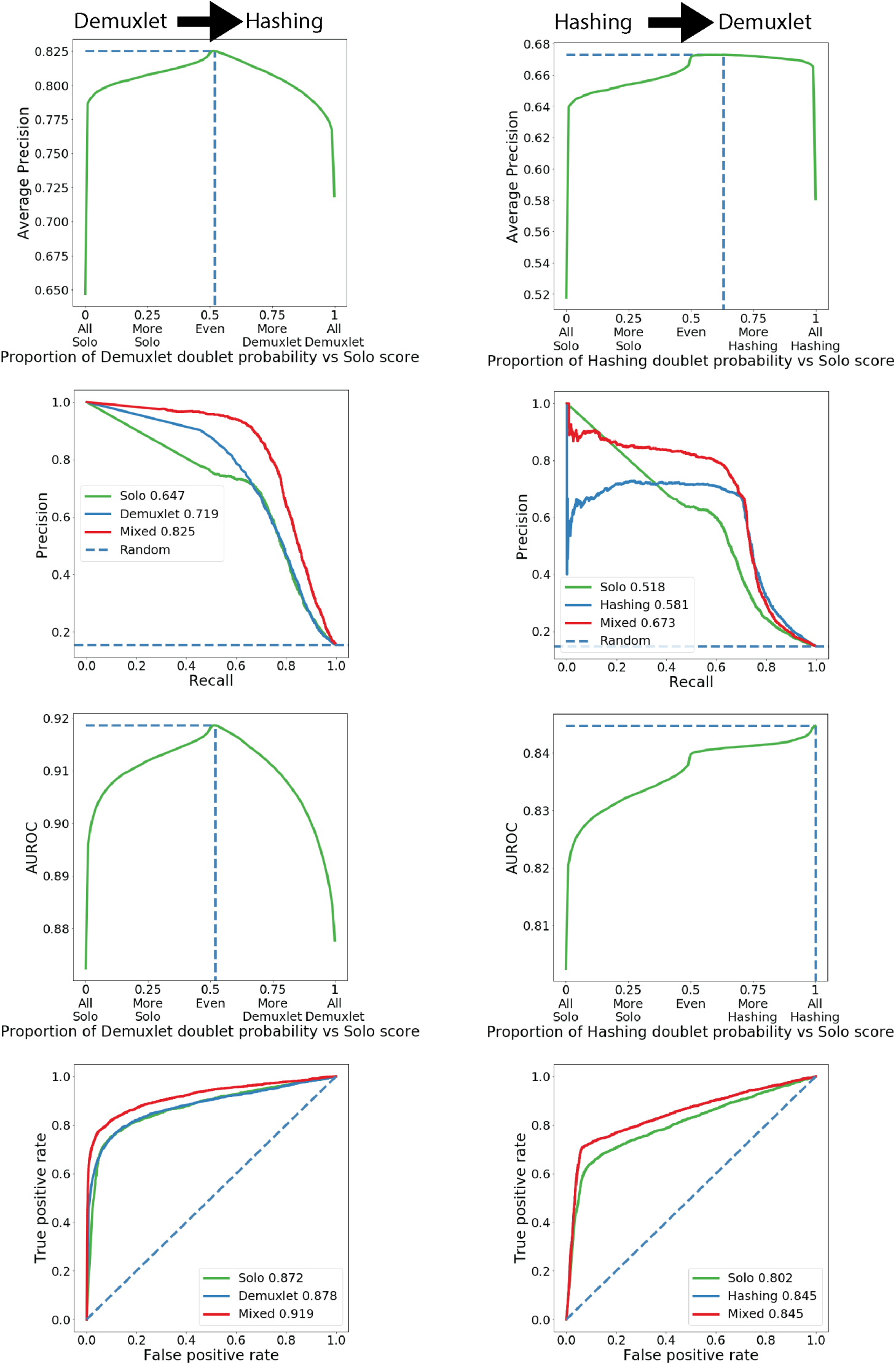
Combining Solo with experimental methods improves doublet classification. We considered doublet identification strategies where both Solo and an experimental method were applied. The Stoeckius et al. HTO8 dataset was generated from PBMCs from eight genetically diverse individuals [10]. Each set of cells was hashed with an antibody conjugated oligo. In the first column, we compute weighted sums of Solo and Demuxlet predictions and compute accuracy with respect to hashing labels. In the second column, we compute weighted sums of Solo and hashing predictions and compute accuracy with respect to Demuxlet labels. Note, that Demuxlet and hashing annotations are not independent because hashes were introduced to each individual’s PBMCs. Thus, the methods will tend to misclassify the same false negatives. This results in a slight bias against Solo in these experiments. In row 1, we plot AP for weighted prediction sums and point to the maximum with a dashed line. In row 2, we plot precision-recall curves for the optimal mixture versus the methods alone. In row 3, we plot AUROC for weighted prediction sums and point to the maximum with a dashed line. In row 4, we plot ROC curves for the optimal mixture versus the methods alone.

**Supplementary Figure 12:**
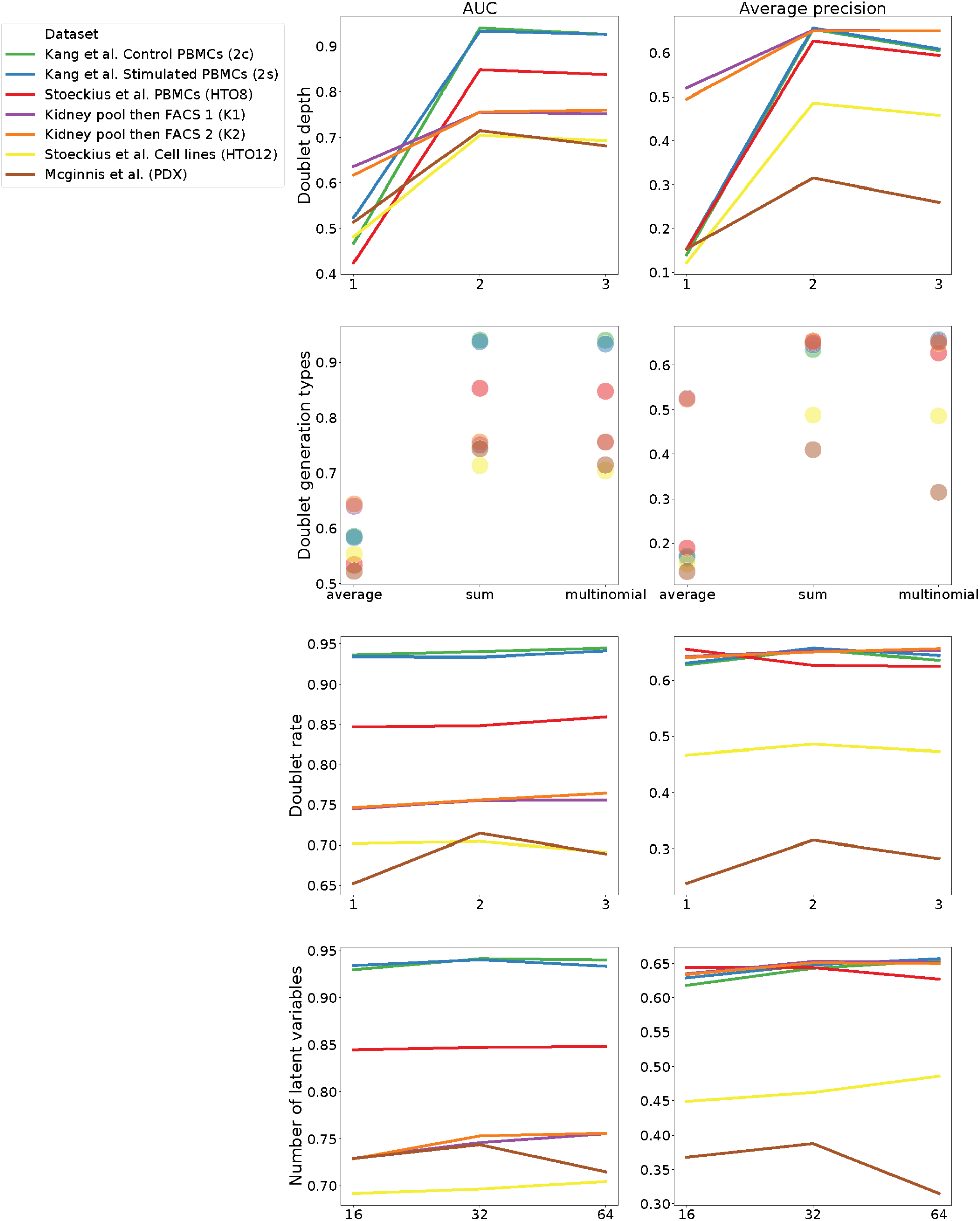
Solo parameter settings. We computed AUROC (left column) and AP (right column) for many different hyperparameter settings. In row 1, we modulated “doublet depth”, which determines the UMI depth for simulated cells by scaling the mean depth of the two sampled cells. I.e. the value two effectively takes the sum of the two cell depths. In row 2, we modulated “doublet generation type”, either taking the exact mean or sum of the two sampled cells or applying our multinomial sampling strategy. In row 3, we modulated “doublet rate”, which determines the number of simulated doublets relative to the observed cells. In row 4, we modulated the number of latent variables in the scVI model.

